## Supplementary Information for "The global diversity of the major parasitic nematode *Haemonchus contortus* is shaped by human intervention and climate"

### Supplementary Data

**Section 1: *Haemonchus contortus* populations are genetically diverse, mediated by large effective population sizes** **3**

Supplementary Table 1. Metadata describing the provenance and sequencing of individual *Haemonchus contortus* samples 3

Supplementary Table 2. Average nucleotide diversity by population 3

Supplementary Figure 1. Observed levels of nucleotide diversity in populations from France, Guadeloupe, and Namibia. 4

**Section 2: Global population connectivity of *Haemonchus contortus* is characterised by old and new migration** **6**

Supplementary Figure 2. Neighbour-joining tree inferred from the pairwise divergence between individual males 6

Supplementary Figure 3. Population clustering by means of a PCA applied to mitochondrial fixed variants derived from consensus sequences 7

Supplementary Figure 4. Pairwise FST estimates binned by MAF between populations with at least 5 individuals 9

Supplementary Table 3. Maximum-likelihood parameter estimates obtained from the joint demographic inference analysis 9

Supplementary Figure 5. Admixture median absolute deviation for K clusters ranging from 2 to 10. 10

Supplementary Figure 6. Admixture pattern across populations for K values of 2, and 4 to 10. 11

Supplementary Figure 7. Bayesian coalescent-based consensus tree of mitochondrial genomes 12

**Section 3: The evolution of anthelmintic resistance has left distinct patterns of diversity on the *Haemonchus contortus* genome** **13**

Supplementary Figure 8. Tajima's *D* estimate plotted against genomic position. 13

Supplementary Figure 9. Reduction of genetic diversity in the vicinity of *Hco-tbb-iso-1* locus for three benzimidazole-resistant and two benzimidazole-susceptible populations. 14

Supplementary Table 4. Population haplotype frequency at SNP in codon 167, 198, 200 of the *Hco-tbb-iso-1* and associated fenbendazole efficacy 15

Supplementary Figure 10. Heatmap of pairwise allelic differences between individual phased genotypes spanning the *Hco-tbb-iso-1* locus. 16

Supplementary Figure 11. Topology weighing analysis of a 100-Kbp window centred on *Hco-tbb-iso-1*. 17

Supplementary Table 5. Genotype counts at each codon position of *Hco-tbb-iso-1* according to the considered GL cut-off 18

Supplementary Figure 12. Mean coverage of genotypic group at SNP in codon positions 167, 198 and 200 of *Hco-btub-1.* 19

Supplementary Figure 13. Individual genotypes at mutant SNP positions of *Hco-tbb-iso1* inferred from genotype likelihoods. 20

Supplementary Figure 14. XP-CLR selection score plotted against genomic position. 21

Supplementary Table 6. Positional candidate genes overlapping significant XP-CLR selection score 21

Supplementary Table 7. Significant GO term enrichment from genes under significant diversifying selection across pairwise comparisons 21

Supplementary Table 8. Significant GO term enrichment from genes under significant diversifying selection in at least one of the population pairwise comparison 21

**Section 4: Climatic adaptation has shaped genomic variation between populations 22**

Supplementary Table 9. Differentiated windows between populations from contrasted climatic conditions 22

Supplementary Table 10. Bioclimatic variables definitions and codes 22

Supplementary Table 11. Significant associations between SNP markers and temperature annual range (BIO7) and annual precipitation (BIO12) bioclimatic variables 22

Supplementary Figure 15. Pairwise Pearson's correlations (a) and principal component analysis (b) between environmental variables from eight populations 23

**Section 5: Supplementary technical notes 24**

Supplementary Figure 16. Variant Quality Score Recalibration (VSQR) summary statistics 24

Supplementary Figure 17. Coverage improvement for a subset of 43 individual *Haemonchus contortus* males 25

Supplementary Figure 18. The relationship between sample coverage and population diversity estimates 26

Supplementary Figure 19. The relationship between pairwise $F_{ST}$ estimates and coverage according to the considered framework 27

Supplementary Figure 20. The relationship between Hamming's distance and coverage 28

Supplementary Figure 21. Principal component analysis based on nuclear VQSR SNP calls across 223 individuals 29

Supplementary Figure 22. Admixture analysis run for chromosome II on the same set of 43 individuals before (left) or after (right) resequencing for K ranging from 2 to 5 30

Supplementary Figure 23. Genotype discordance after Beagle imputation 32

**Supplementary references 33**

#### Section 1: *Haemonchus contortus* populations are genetically diverse, mediated by large effective population sizes

##### Supplementary Table 1. Metadata describing the provenance and sequencing of individual *Haemonchus contortus* samples

- see accompanying file

Unless stated otherwise (France, Guadeloupe, South-Africa and Australia), isolates were sampled from a single farm within each country.

##### Supplementary Table 2. Average nucleotide diversity by population

| Population | Mean π across nuclear genome | Standard deviation | Population mean coverage |
|---:|---:|---:|---:|
| ACO | 0.00681 | 0.00096 | 1.79 |
| AUS.1 | 0.00613 | 0.00044 | 2.57 |
| AUS.2 | 0.00509 | 0.00032 | 2.74 |
| BEN | 0.00939 | 0.00232 | 0.33 |
| CAP | 0.00721 | 0.00104 | 0.70 |
| FRA.1 | 0.00723 | 0.00037 | 3.32 |
| FRA.2 | 0.00544 | 0.00059 | 1.82 |
| FRA.3 | 0.00623 | 0.00084 | 0.48 |
| FRG | 0.01078 | 0.00032 | 4.40 |
| IND | 0.00555 | 0.00075 | 4.95 |
| MOR | 0.00820 | 0.00044 | 3.50 |
| NAM | 0.01300 | 0.00075 | 5.62 |
| STA.1 | 0.00669 | 0.00183 | 4.01 |
| STA.2 | 0.00442 | 0.00045 | 1.79 |
| STA.3 | 0.01124 | 0.00090 | 2.64 |
| STO | 0.00975 | 0.00075 | 2.61 |
| ZAI | 0.00858 | 0.00029 | 2.11 |
| FRA.1* | 0.00610 | 0.00049 | 8.03 |
| FRG* | 0.01063 | 0.00079 | 12.75 |
| NAM* | 0.01136 | 0.00079 | 9.85 |

Asterisks indicates population subset of 5 individuals with minimum mean coverage of 5×: this was limited to France (FRA.1, n = 5, mean coverage of 7.66×), Guadeloupe (FRG, n = 5, mean coverage of 12.75×) and Namibia (NAM, n = 6, mean coverage of 9.85×).

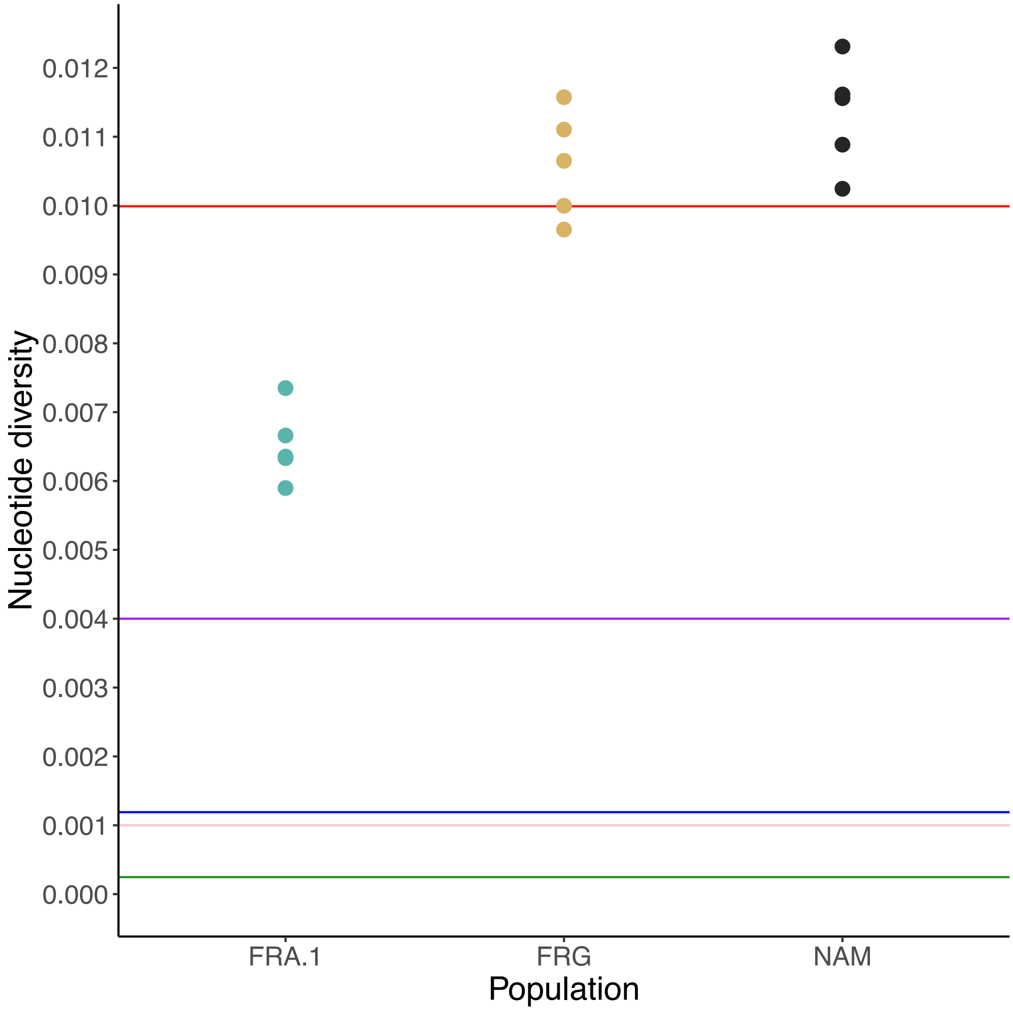


**Supplementary Figure 1. Observed levels of nucleotide diversity in populations from France, Guadeloupe, and Namibia.**

Nucleotide diversity (π) is presented representing the number of substitutions per base for three subsets of populations built from individuals with a mean coverage of 5x and more, i.e. France (FRA.1, n = 5, mean coverage of 7.66×), Guadeloupe (FRG, n = 5, mean coverage of 12.75×) and Namibia (NAM, n = 6, mean coverage of 9.85×). Each dot corresponds to the average nucleotide diversity across single autosome. Horizontal lines provide genome-wide nucleotide diversity estimates from two other parasitic nematodes (Onchocercidae, clade III): *Wuchereria bancrofti*[1] ($\pi = 2.7 \times 10^{-4}$, green), and *Onchocerca volvulus*[2] ($\pi_S = 4 \times 10^{-3}$ and $\pi_N = 1 \times 10^{-3}$ in purple and pink respectively). Red horizontal line matches previously reported values for *Drosophila melanogaster*[3] (0.999% averaged across three populations showing values of 0.00531, 0.00752, 0.01714).

#### Section 2: Global population connectivity of *Haemonchus contortus* is characterised by old and new migration

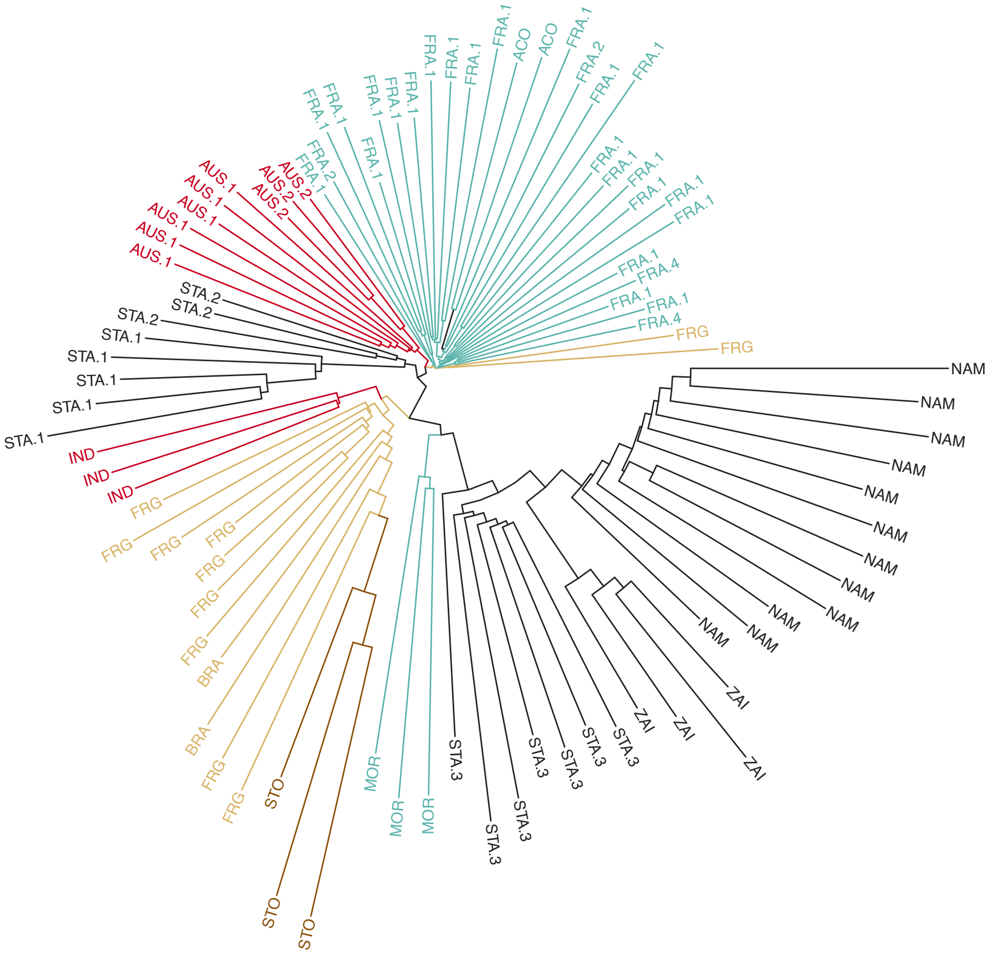


**Supplementary Figure 2. Neighbour-joining tree inferred from the pairwise divergence between individual males**

Pair-wise Hamming's distance, *i.e.* the SNP fraction non-identical-by-state between two individuals, were computed with PLINK[6] v1.90b3v. Because of coverage bias, the analysis was restricted to 75 individuals with a minimum mean coverage of 2.5x.

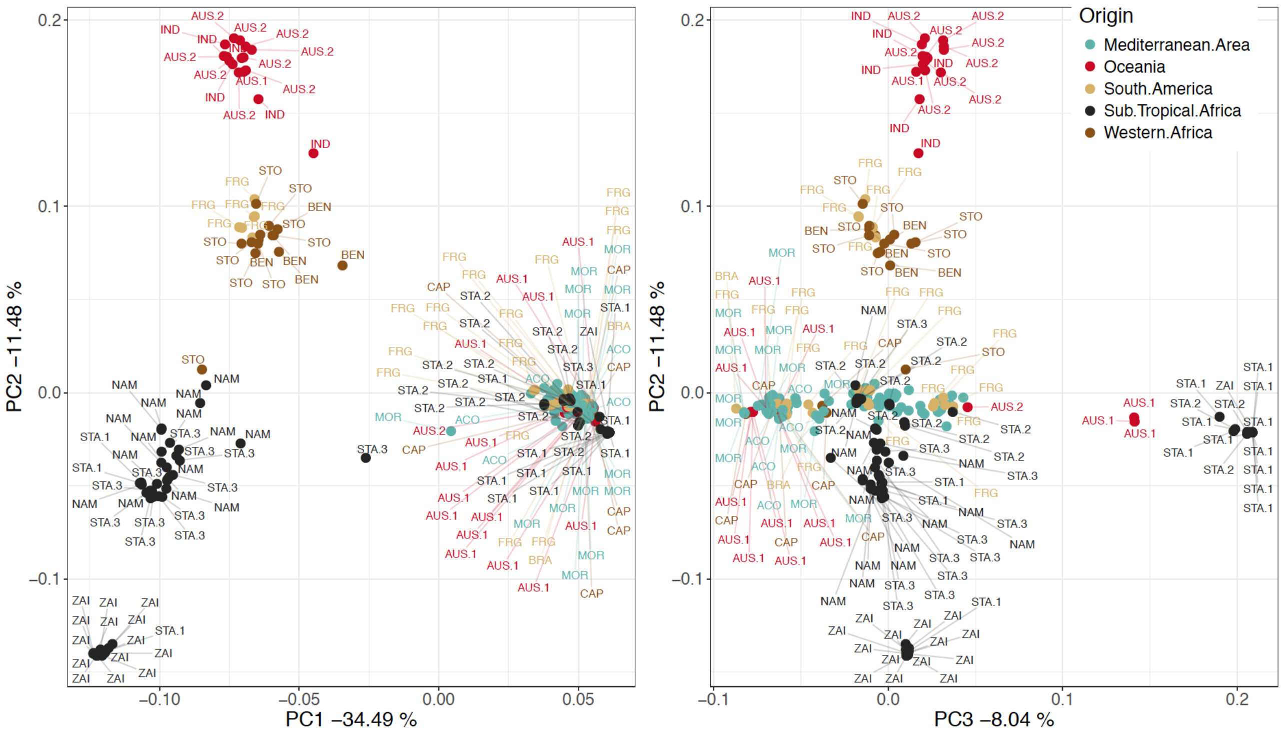


**Supplementary Figure 3. Population clustering by means of a PCA applied to mitochondrial fixed variants derived from consensus sequences**

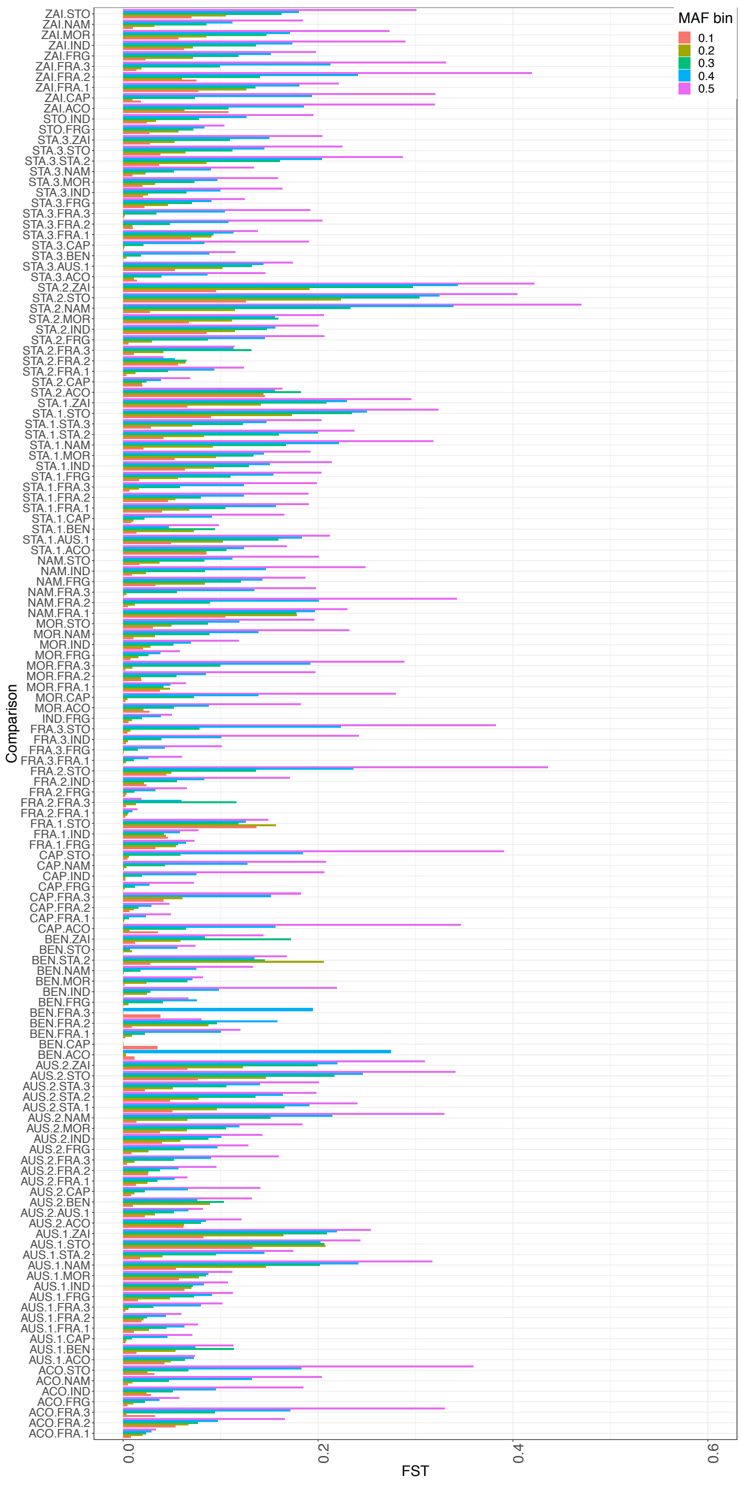

MAF bin
0.1
0.2
0.3
0.4
0.5
Comparison
ZAI.STO
ZAI.NAM
ZAI.MOR
ZAI.IND
ZAI.FRG
ZAI.FRA.3
ZAI.FRA.2
ZAI.FRA.1
ZAI.CAP
ZAI.ACO
STO.IND
STO.FRG
STA.3.ZAI
STA.3.STO
STA.3.STA.2
STA.3.NAM
STA.3.MOR
STA.3.IND
STA.3.FRG
STA.3.FRA.3
STA.3.FRA.2
STA.3.FRA.1
STA.3.CAP
STA.3.BEN
STA.3.AUS.1
STA.3.ACO
STA.2.ZAI
STA.2.STO
STA.2.NAM
STA.2.MOR
STA.2.IND
STA.2.FRG
STA.2.FRA.3
STA.2.FRA.2
STA.2.FRA.1
STA.2.CAP
STA.2.ACO
STA.1.ZAI
STA.1.STO
STA.1.STA.3
STA.1.STA.2
STA.1.NAM
STA.1.MOR
STA.1.IND
STA.1.FRG
STA.1.FRA.3
STA.1.FRA.2
STA.1.FRA.1
STA.1.CAP
STA.1.BEN
STA.1.AUS.1
STA.1.ACO
NAM.STO
NAM.IND
NAM.FRG
NAM.FRA.3
NAM.FRA.2
NAM.FRA.1
MOR.STO
MOR.NAM
MOR.IND
MOR.FRG
MOR.FRA.3
MOR.FRA.2
MOR.FRA.1
MOR.CAP
MOR.ACO
IND.FRG
FRA.3.STO
FRA.3.IND
FRA.3.FRG
FRA.3.FRA.1
FRA.2.STO
FRA.2.IND
FRA.2.FRG
FRA.2.FRA.3
FRA.2.FRA.1
FRA.1.STO
FRA.1.IND
FRA.1.FRG
CAP.STO
CAP.NAM
CAP.IND
CAP.FRG
CAP.FRA.3
CAP.FRA.2
CAP.FRA.1
CAP.ACO
BEN.ZAI
BEN.STO
BEN.STA.2
BEN.NAM
BEN.MOR
BEN.IND
BEN.FRG
BEN.FRA.3
BEN.FRA.2
BEN.FRA.1
BEN.CAP
BEN.ACO
AUS.2.ZAI
AUS.2.STO
AUS.2.STA.3
AUS.2.STA.2
AUS.2.STA.1
AUS.2.NAM
AUS.2.MOR
AUS.2.IND
AUS.2.FRG
AUS.2.FRA.3
AUS.2.FRA.2
AUS.2.FRA.1
AUS.2.CAP
AUS.2.BEN
AUS.2.AUS.1
AUS.2.ACO
AUS.1.ZAI
AUS.1.STO
AUS.1.STA.2
AUS.1.NAM
AUS.1.MOR
AUS.1.IND
AUS.1.FRG
AUS.1.FRA.3
AUS.1.FRA.2
AUS.1.FRA.1
AUS.1.CAP
AUS.1.BEN
AUS.1.ACO
ACO.STO
ACO.NAM
ACO.IND
ACO.FRG
ACO.FRA.3
ACO.FRA.2
ACO.FRA.1
0.0
0.2
0.4
0.6
FST

**Supplementary Figure 4. Pairwise FST estimates binned by MAF between populations with at least 5 individuals**

**Supplementary Table 3. Maximum-likelihood parameter estimates obtained from the joint demographic inference analysis**

- see accompanying file

For every population, unscaled parameters from forward genetic simulations run in δaδi are listed. Initial exploration of the fit of more complex scenarios than a simple split and isolation model suggested there was little power in our data to estimate parameters accurately under these models. Nevertheless, outputs from these models are shown as an indication of the most likely demographic scenario between corresponding populations.

For each model and population couple (population1-population2), estimation round and replicate are provided along with considered genomic segment length (L), model log-likelihood, Akaike Information Criterion (AIC). Nu1a and nu2a correspond to population sizes after first event and nu1b and nu2b indicate population size following secondary event if any. In case of asymmetrical migration between populations, two migration rates are provided (m12, m21 for migration from population1 to population2 and from population2 to population 1 respectively). Unscaled timings of events appear as T1 (time of split), T2 (time since secondary event) and T3 (time since third event if any) and corresponding years under the common era (CE) have been listed as Year of split, Year of secondary event, Year of isolation respectively. Standard deviation of parameters estimates are indicated in brackets for simple "split and isolation" models. Asym: asymmetrical; Sym: symmetrical.

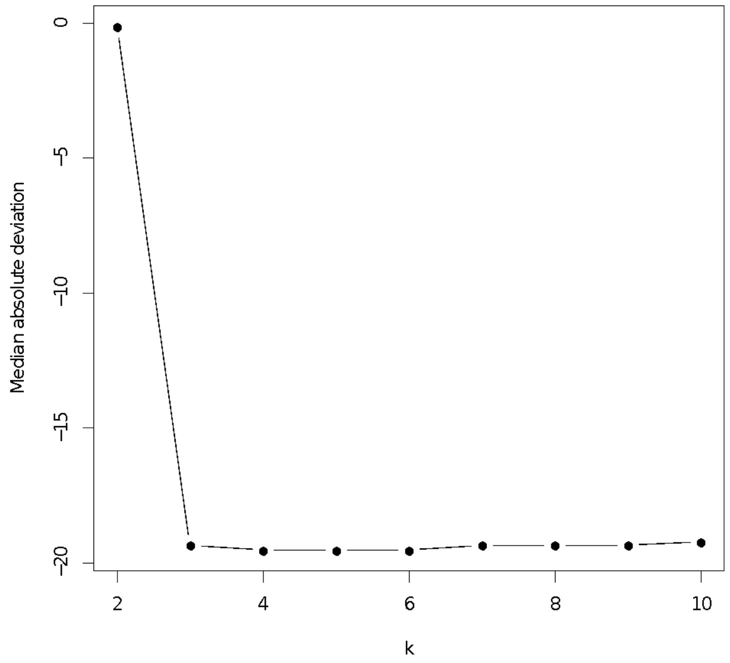


**Supplementary Figure 5. Admixture median absolute deviation for K clusters ranging from 2 to 10.**

Median absolute deviation was estimated across five runs of NGSAdmix[7], retaining sites with less than 50% missing data across individuals and minor allele frequency (MAF) above 5% and leaving one autosome out at a time.

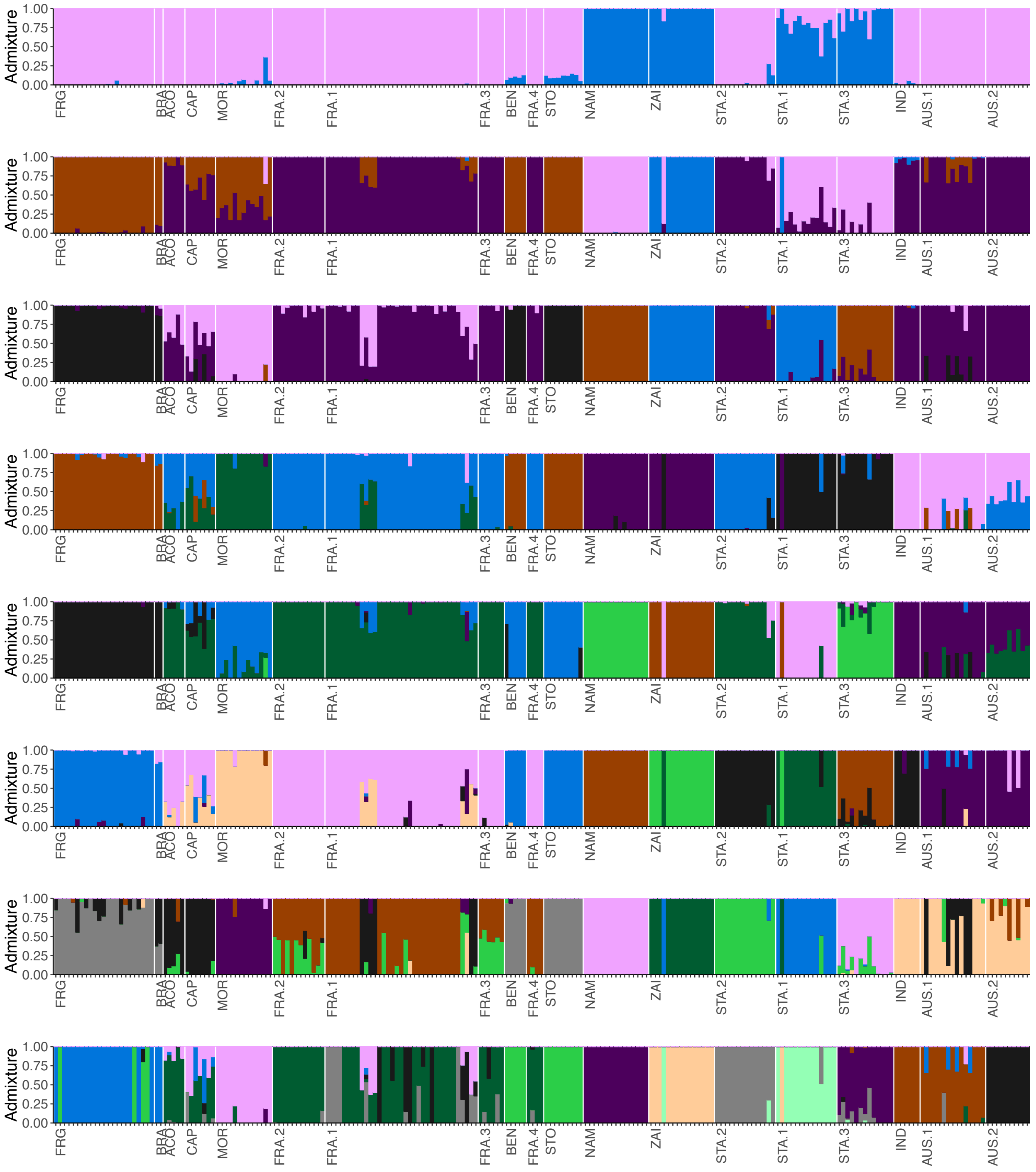


**Supplementary Figure 6. Admixture pattern across populations for K values of 2, and 4 to 10.**

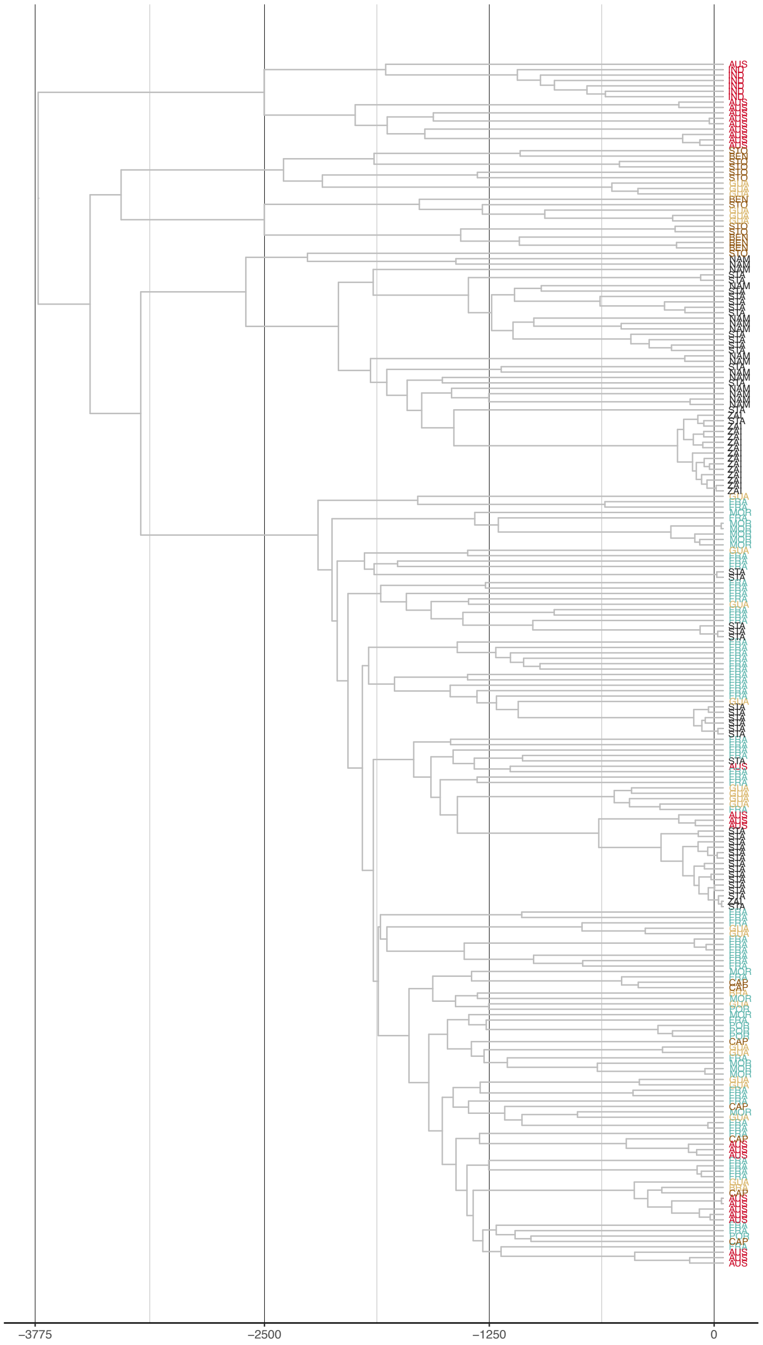


**Supplementary Figure 7. Bayesian coalescent-based consensus tree of mitochondrial genomes**

Maximum clade credibility tree is represented from 50 million Monte-Carlo Markov Chains, and burn-in of the first 20 million iterations. Tips are coloured according to the sample geographical location following the same key as in Fig.1. Vertical lines correspond to 1250-year intervals.

#### Section 3: The evolution of anthelmintic resistance has left distinct patterns of diversity on the *Haemonchus contortus* genome

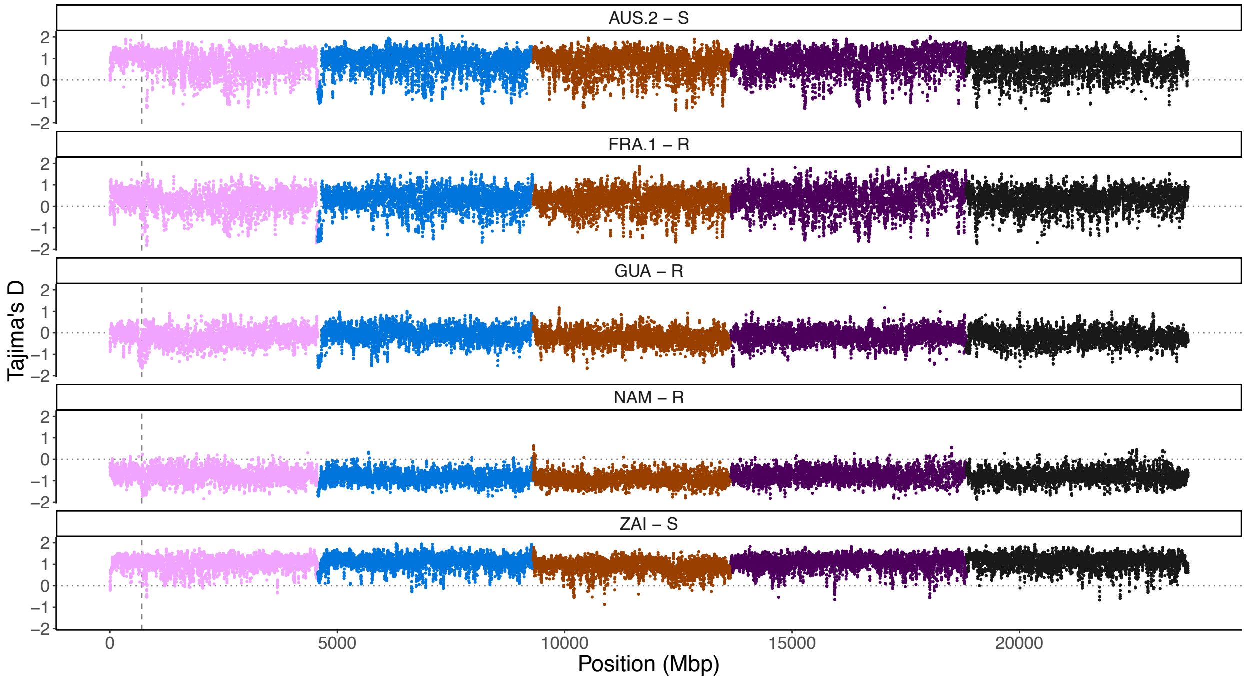


**Supplementary Figure 8. Tajima's *D* estimate plotted against genomic position.**

Plot represents Tajima's *D* coefficient along the genome for five populations with the highest representation of individuals with a mean depth of coverage higher than 5x. Vertical dash line indicates β-tubulin locus.

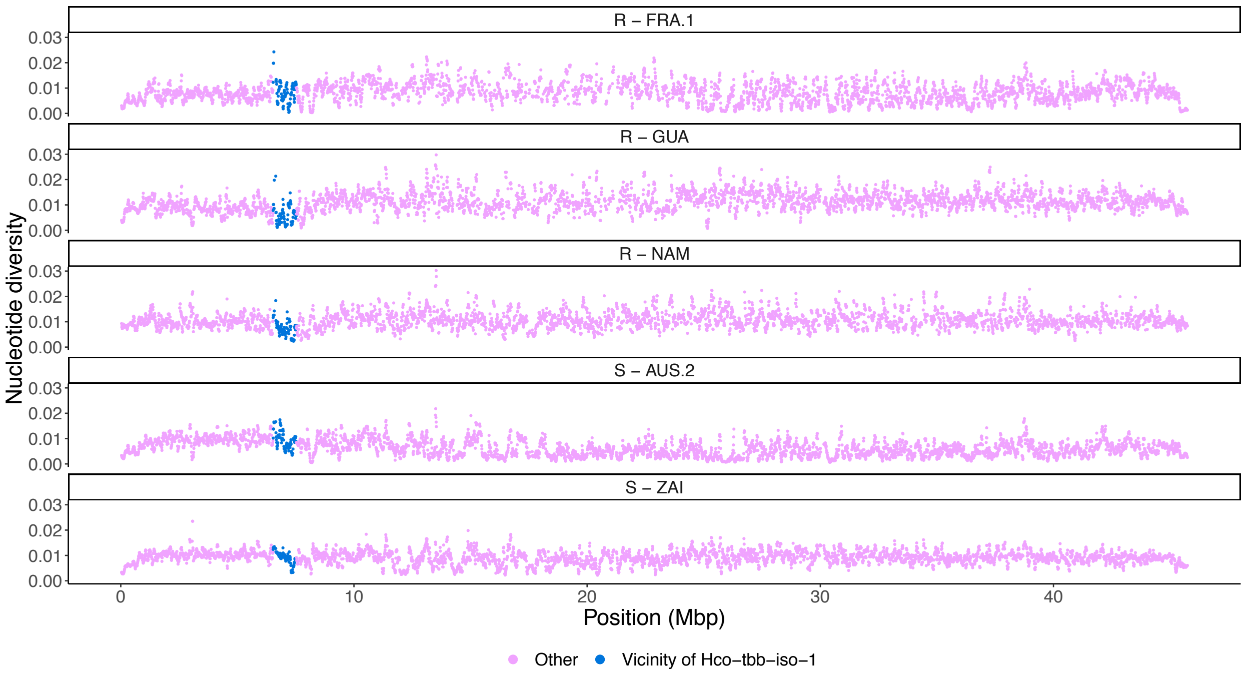


**Supplementary Figure 9. Reduction of genetic diversity in the vicinity of *Hco-tbb-iso-1* locus for three benzimidazole-resistant and two benzimidazole-susceptible populations.** Plot represents nucleotide diversity along chromosome I for five populations with the highest representation of individuals with a mean depth of coverage higher than 5x (R: resistant; S: susceptible). Blue dots highlight the diversity reduction within a 1-Mbp window centred at the *β*-tubulin locus (mean π = 0.007 across resistant populations) relative to chromosome I (mean π = 0.011 across resistant populations). Diversity in susceptible populations was slightly higher at the *β*-tubulin locus (mean π = 0.009 across resistant populations) relative to chromosome I (mean π = 0.008 across resistant populations). Direct comparison between resistant and susceptible populations was not possible because of the difference in mean coverage.

**Supplementary Table 4. Population haplotype frequency at SNP in codon 167, 198, 200 of the *Hco-tbb-iso-1* and associated fenbendazole efficacy**

.

| Population | A/A-A/A-T/T | A/T-A/A-T/T | T/T-A/A-A/A | T/T-A/A-T/A | **T/T-A/A-T/T** | T/T-C/A-A/T | T/T-C/C-T/T | Fenbendazole efficacy |
|---|---|---|---|---|---|---|---|---|
| ACO | 0 | 0 | 0 | 0 | 1 | 0 | 0 | N/A |
| AUS.1 | 0 | 0 | 5 | 0 | 0 | 1 | 2 | 43 |
| AUS.2 | 0 | 0 | 0 | 0 | 4 | 0 | 0 | 100 |
| BRA | 0 | 0 | 1 | 0 | 0 | 0 | 0 | 0.795 μg/ml[a] |
| CAP | 0 | 0 | 0 | 0 | 1 | 0 | 0 | N/A |
| FRA.1 - Farm 1 | 0 | 0 | 2 | 0 | 0 | 0 | 0 | 60.2 |
| FRA.1 - Farm 2 | 0 | 0 | 0 | 0 | 0 | 0 | 4 | 64.3 |
| FRA.1 - Farm 3 | 0 | 0 | 2 | 0 | 0 | 0 | 0 | N/A |
| FRA.1 - Farm 4 | 0 | 0 | 1 | 0 | 0 | 1 | 0 | 50.4 |
| FRA.1 - Farm 5 | 1 | 1 | 0 | 0 | 0 | 0 | 0 | 98.6 |
| FRA.1 - Farm 6 | 0 | 0 | 0 | 0 | 0 | 0 | 2 | 60.4 |
| FRA.1 - Farm 8 | 0 | 0 | 2 | 0 | 0 | 0 | 0 | 43.7 |
| FRA.2 | 0 | 0 | 0 | 0 | 3 | 0 | 0 | 100 |
| FRA.4 | 0 | 0 | 0 | 0 | 1 | 0 | 0 | 100 |
| GUA | 0 | 0 | 14 | 0 | 0 | 0 | 0 | N/A |
| IND | 0 | 0 | 0 | 0 | 1 | 0 | 0 | N/A |
| MOR | 0 | 0 | 1 | 0 | 1 | 0 | 0 | N/A |
| NAM | 0 | 0 | 4 | 1 | 0 | 0 | 0 | N/A |
| STA.1 | 0 | 0 | 1 | 0 | 1 | 1 | 2 | 0.587 μg/ml[a] |
| STA.2 | 0 | 0 | 0 | 0 | 2 | 0 | 0 | N/A |
| STA.3 | 0 | 0 | 6 | 0 | 0 | 0 | 0 | 33.75[b] |
| STO | 0 | 0 | 0 | 0 | 3 | 0 | 0 | N/A |
| ZAI | 0 | 0 | 0 | 0 | 1 | 0 | 0 | 100 |

For each population, the total count of each genotype combination at codon positions 167/198/200 is listed (based on individuals showing minimum genotype likelihood of 60%). Fully susceptible reference genotype is T/T-A/A-T/T for codon positions 167 -198 - 200. Any variation in this sequence indicates the recessive mutant. Unless stated otherwise, benzimidazole efficacy refers to Faecal Egg Count Reduction Test (FECRT, in %) value after fenbendazole treatment.

a: Median inhibitory concentration (IC50) for triclabendazole measured by means of an egg hatch assay and relative to a susceptible isolate showing an IC50 of 0.022 μg/ml.

b: FECRT value after albendazole treatment[8].

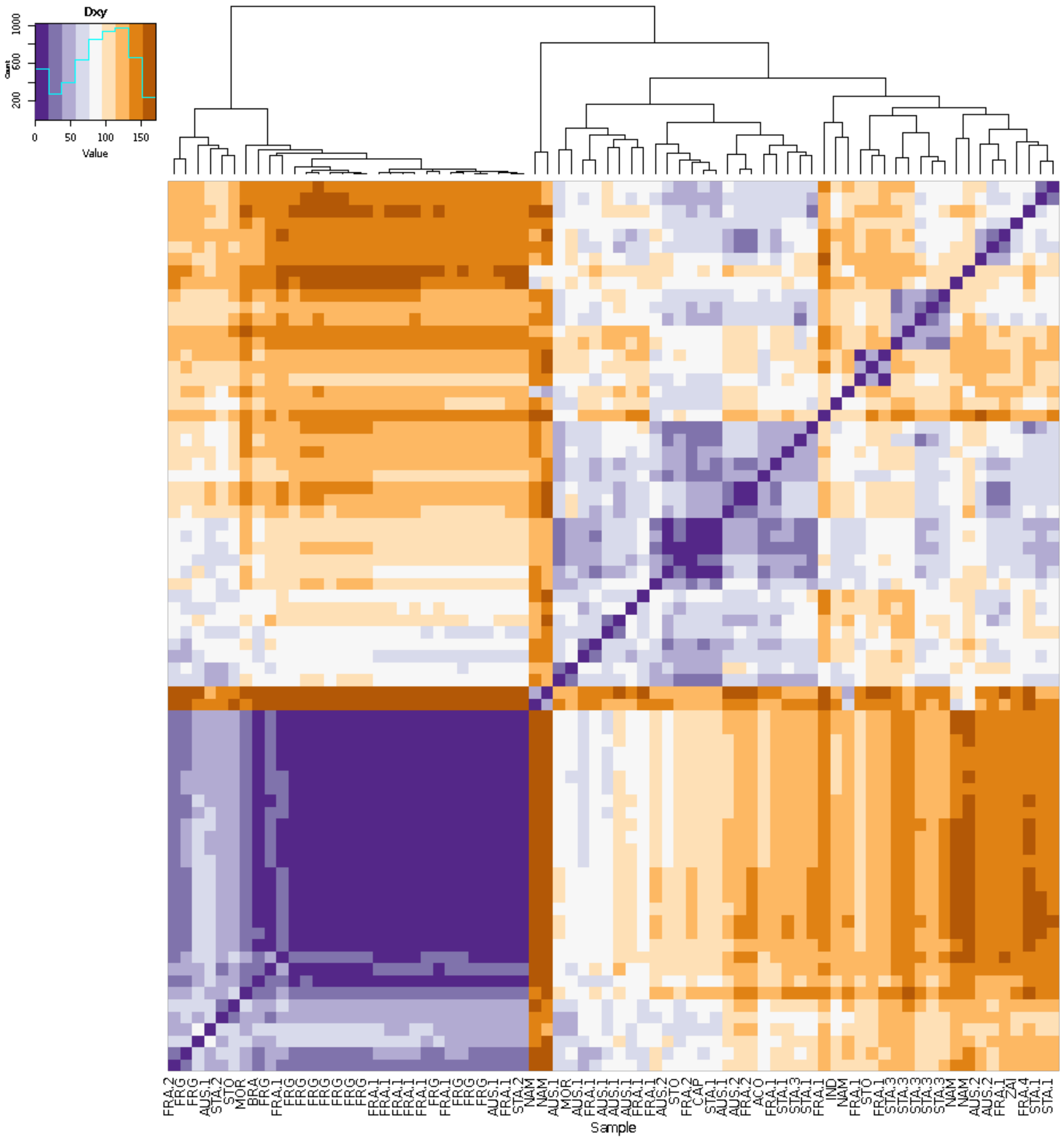


**Supplementary Figure 10. Heatmap of pairwise allelic differences between individual phased genotypes spanning the *Hco-tbb-iso-1* locus.**

Number of pairwise allelic differences between phased genotypes over the *Hco-tbb-iso1* locus (2,158 bp, 296 SNP positions considered) are plotted for the 74 individuals with minimal genotype likelihood of 60%. Individual clustering reveals close sequence similarities (purple) between FRA.1 and GUA individuals, whereas other samples tend to cluster in the same way as predicted from the mitochondrial and nuclear genomes.

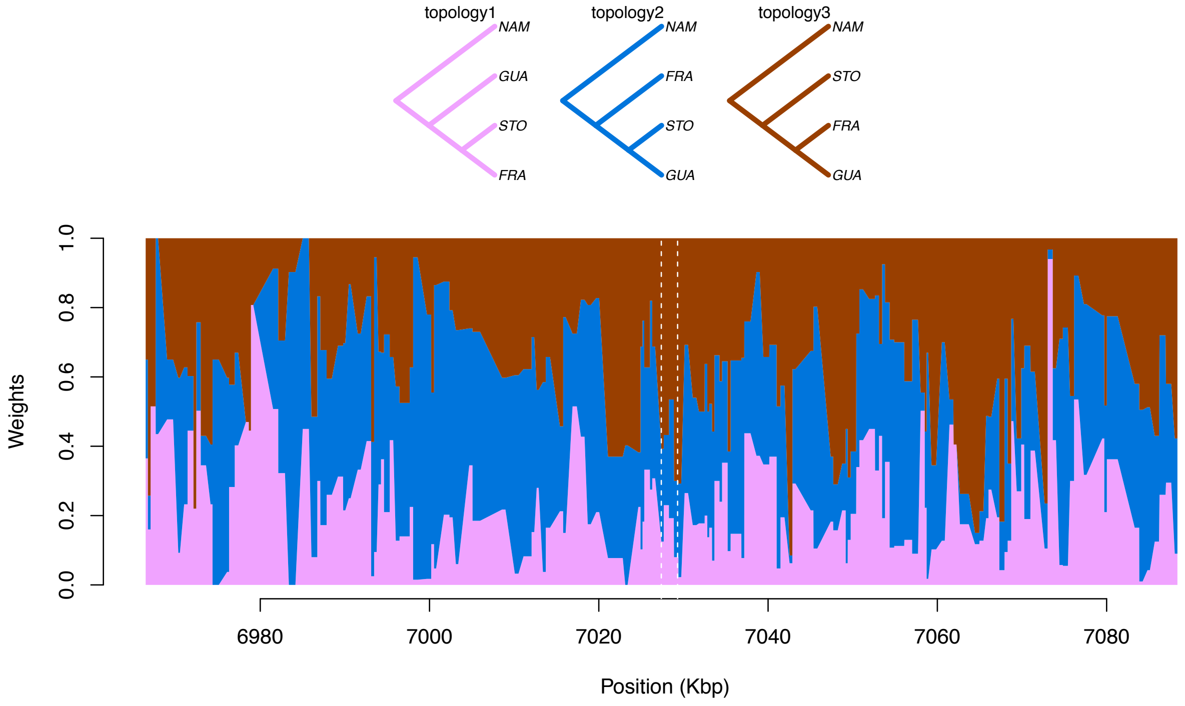


**Supplementary Figure 11. Topology weighing analysis of a 100-Kbp window centred on *Hco-tbb-iso-1*.**

Topology weighing analysis of a 20-Kbp window centred on *Hco-tbb-iso-1*, using populations from Namibia (NAM), France (FRA.1), Guadeloupe (FRG) and São Tomé (STO). At each position, the weight of each of the three possible topologies inferred from 50 Kbp-windows is overlaid. Topology 2 (blue) corresponds to an isolation-by-distance history, while topology 3 (brown) would agree with shared introgressed material between worm populations from French mainland into Guadeloupe. The position of *Hco-tbb-iso-1* locus is indicated by the vertical dashed lines. This figure supports the introgression event and the analysis using Moroccan worms as a link population with FRG.

Genotypes for the *Hco-tbb-iso-1* locus were determined for each of the 223 samples using ANGSD[9]. Analysis of all genotypes prior to filtering (GQ>0) revealed an over-representation of susceptible genotypes at each position (Supplementary Table 3).

**Supplementary Table 5. Genotype counts at each codon position of *Hco-tbb-iso-1* according to the considered GL cut-off**

| | | | P167Y | | | E198A | | | P200Y |
|---|---|---|---|---|---|---|---|---|---|
| | A/A | T/A | **T/T** | **A/A** | A/C | C/C | A/A | T/A | **T/T** |
| GL>0 | 1 | 1 | **221** | **208** | 3 | 12 | 55 | 5 | **166** |
| GL>0.6 | 1 | 1 | **72** | **61** | 3 | 10 | 39 | 4 | **31** |
| GL>0.8 | 0 | 0 | **31** | **25** | 3 | 3 | 17 | 4 | **10** |

*Reference genotypes are indicated in bold.*

A regression analysis of genotype counts at position 200 upon sample mean coverage demonstrated a significant bias ($P$ = 6.64 x $10^{-8}$, $F_{2,220}$ = 17.83) toward reference susceptible genotype T/T in samples with lower coverage (-0.8x for this particular subpopulation). The same applied for position 198 (Kruskal-Wallis $\chi^2$ = 35.423, *df* = 2, $P$ = 0.05).

This bias was corrected for by selecting the only genotype with GL above 60%, resulting in 0.04x difference between homozygous genotypes (Kruskal-Wallis $\chi^2$ = 3.11, *df* =2, $P$ = 0.2115 and $\chi^2$ = 3.74, *df* =2, $P$ = 0.15 for the P200Y and E198A positions respectively; Supplementary Fig. 9). Samples with the reference genotype at codon position 200, still displayed a lower coverage than their counterparts. Applying a more stringent GL cut-off resulted in the same pattern, ruling out a coverage bias as population coverage reached an average of 6x [ 2 – 15.24] in this case. Genotype prediction corroborated known fenbendazole resistance status of Australian (AUS.1) and South-African (STA.3) populations (Supplementary Fig. 9) and were also in line for the French population despite the low number of observations available (Supplementary Table 4).

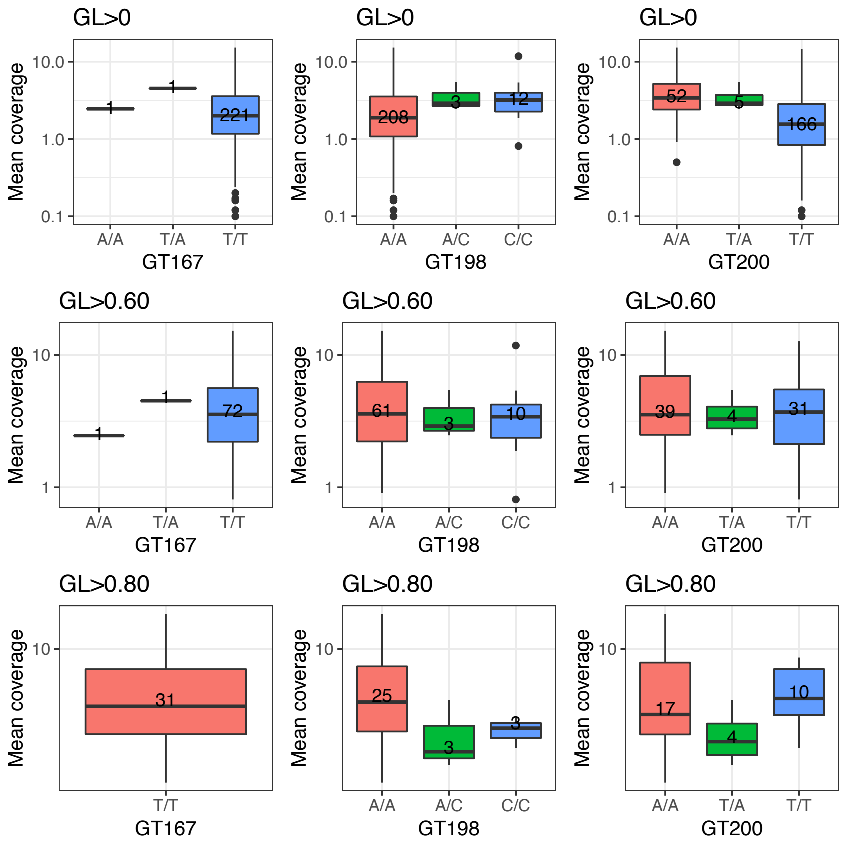


**Supplementary Figure 12. Mean coverage of genotypic group at SNP in codon positions 167, 198 and 200 of *Hco-btub-1.***

For each codon position (GT167, 198 and 200) and genotype likelihood (GL) cut-off, genotypic group mean coverage is represented, showing a coverage bias in reference genotypic group when no GL filtering is applied. No significant impact on the relationship between predicted genotype and samples mean coverage is observed between GL cut-off of 0.6 or 0.8. Number of observations per group are indicated within each box.

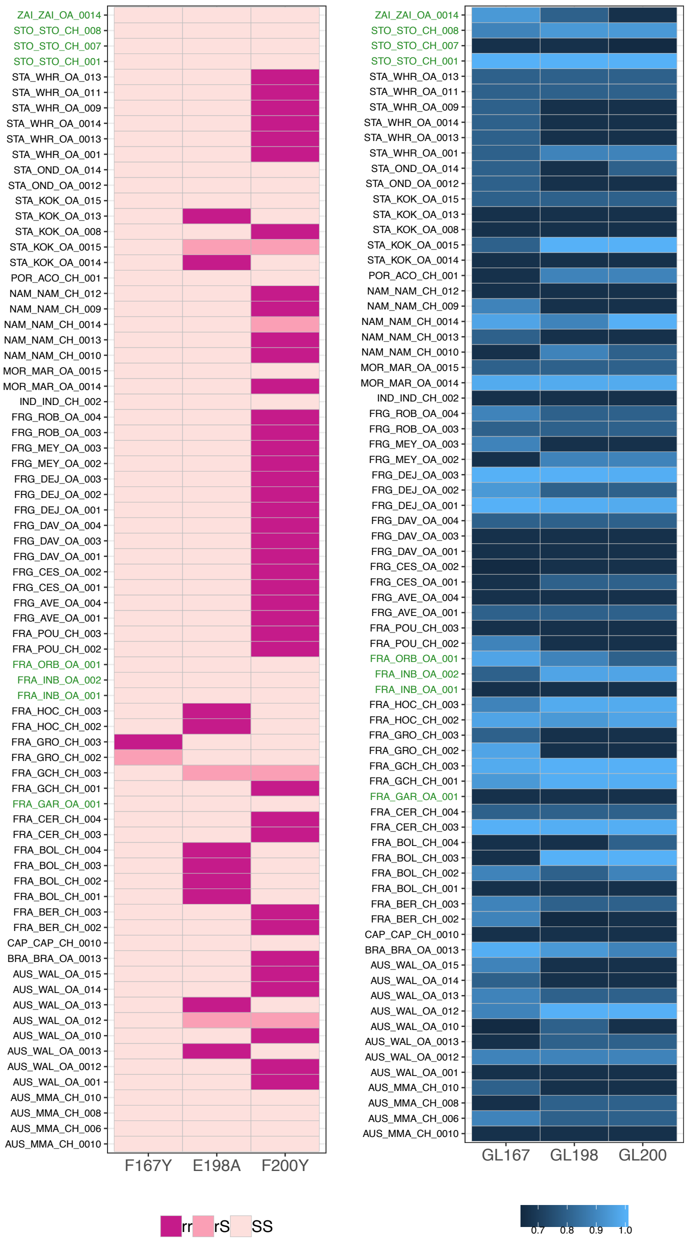


**Supplementary Figure 13. Individual genotypes at mutant SNP positions of *Hco-tbb-iso1* inferred from genotype likelihoods.**

The left panel displays the individual (row) combination of the genotypes at codon position 167, 198 and 200 (column) of the *Hco-tbb-iso-1* locus. Colour intensity follows the mutant allele dosage, i.e. purple standing for mutant homozygotes (rr), salmon for heterozygotes and pink shows homozygote reference genotype. Populations with phenotypically susceptible are coloured in green. The right panel represents the genotype likelihood of each individual-locus combination, ranging from 60% to 100% probability, and showing the absence of bias between coverage and genotype.

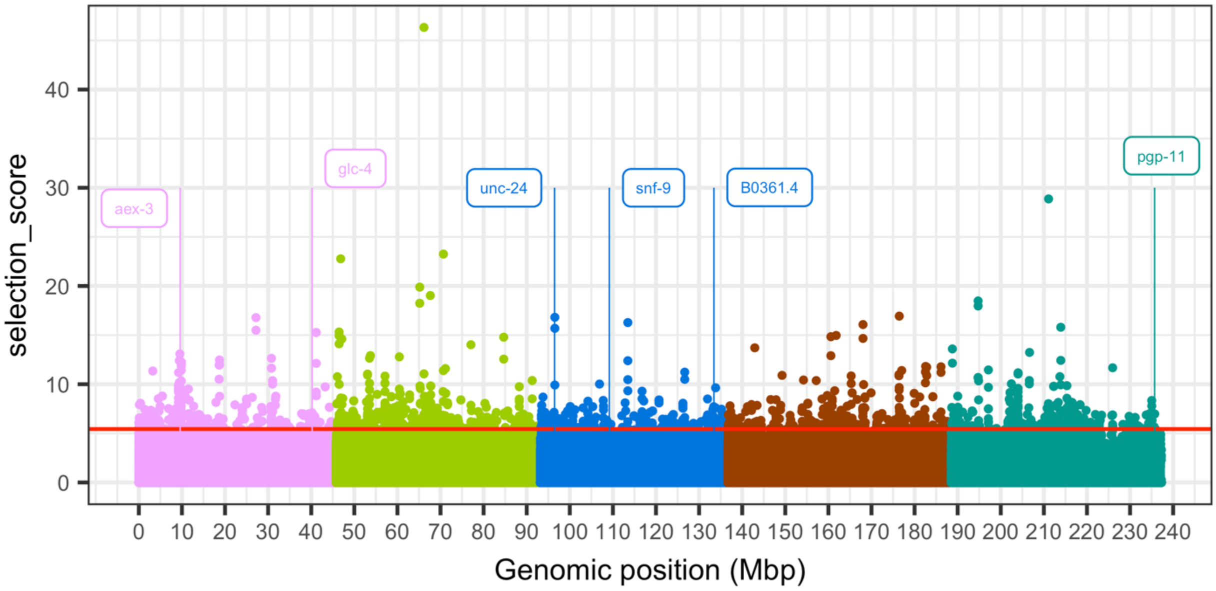


**Supplementary Figure 14. XP-CLR selection score plotted against genomic position.**

Each point represents the selection score generated by XP-CLR by its position in the genome. Positions are given in Mbp, and points are coloured by chromosome. The vertical grey dashed line indicates the *Hco-tbb-iso-1* locus, and the horizontal red line corresponds to the top 0.1% quantile. Coloured vertical lines and boxes point at candidate genes of interest, being either members of the dauer pathway (*daf-36, tax-4*), or putative candidates underpinning ivermectin resistance.

**Supplementary Table 6. Positional candidate genes overlapping significant XP-CLR selection score**

- see accompanying file

**Supplementary Table 7. Significant GO term enrichment from genes under significant diversifying selection across pairwise comparisons**

- see accompanying file

**Supplementary Table 8. Significant GO term enrichment from genes under significant diversifying selection in at least one of the population pairwise comparison**

- see accompanying file

#### Section 4: Climatic adaptation has shaped genomic variation between populations

**Supplementary Table 9. Differentiated windows between populations from contrasted climatic conditions**

- see accompanying file

**Supplementary Table 10. Bioclimatic variables definitions and codes**

- see accompanying file

**Supplementary Table 11. Significant associations between SNP markers and temperature annual range (BIO7) and annual precipitation (BIO12) bioclimatic variables**

- see accompanying file

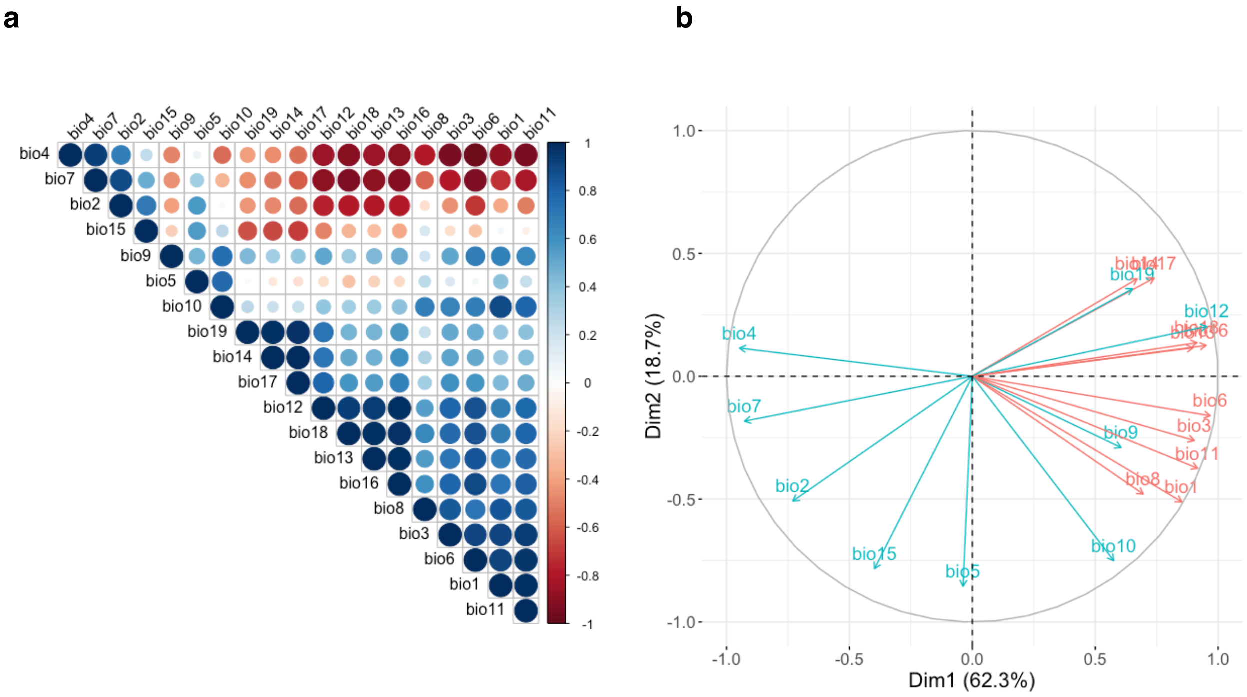


**Supplementary Figure 15. Pairwise Pearson's correlations (a) and principal component analysis (b) between environmental variables from eight populations**

(a) Matrix shows Pearson's correlation between bioclimatic variables determined from GPS coordinates for eight populations. Circle sizes indicate correlation intensity and colors correspond to correlation direction. Variables are clustered according to the correlation they entertain.

(b) Correlation circle shows bioclimatic variables coordinates on first two principal components. Red variables were considered as highly correlated and not considered further for gradient forest analysis.

#### Section 5: Supplementary technical notes

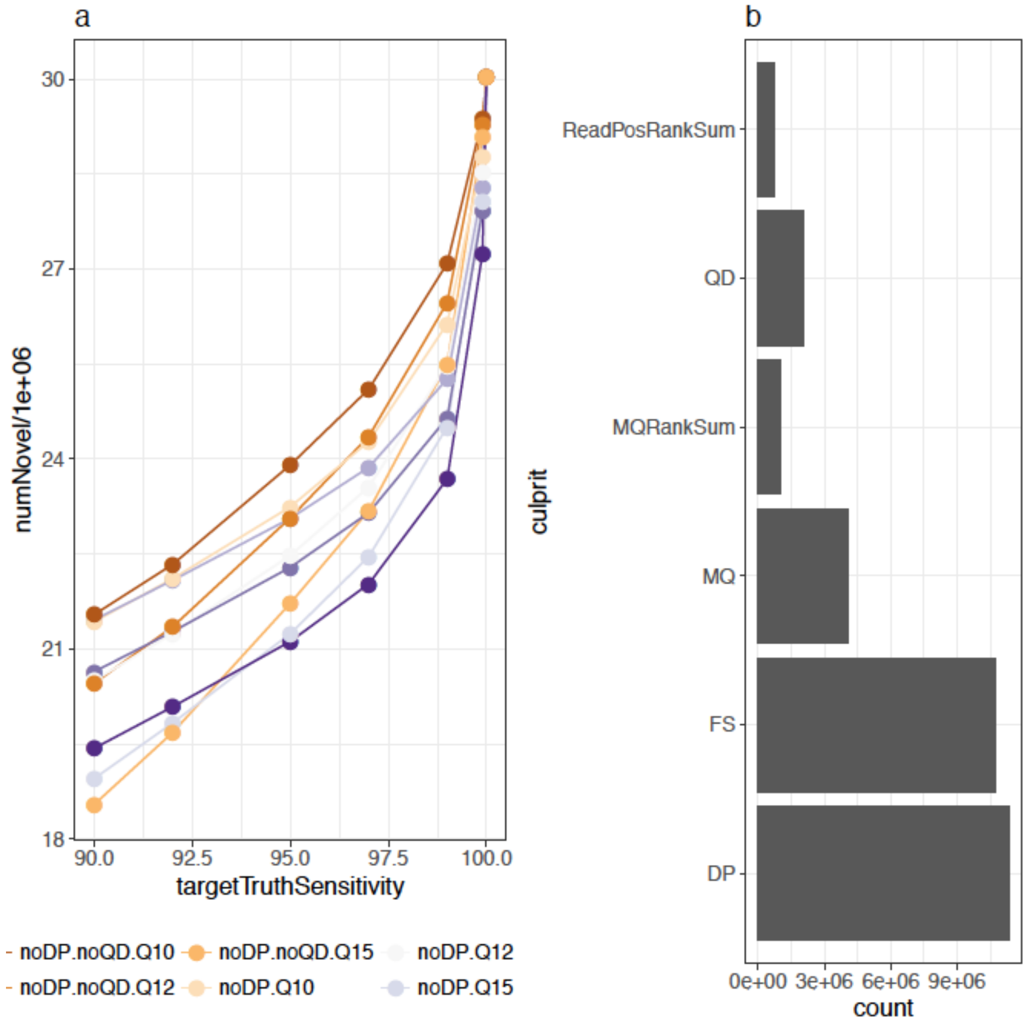


**Supplementary Figure 16. Variant Quality Score Recalibration (VSQR) summary statistics**

##### ***Evaluation of genotype analysis frameworks in face of low coverage samples***

Our sequencing effort achieved a mean coverage of 3x across 223 individuals, thus hampering stringent filtering on SNP genotypes. Two options were implemented to deal with this matter: (i) the Variant Quality Score Recalibration of the GATK[10] software, or (ii) the probabilistic framework implemented in the ANGSD[9] software that relies on genotype likelihoods (GLs). VQSR SNP calls were considered for analyses that were not available in ANGSD, or when bias was identified in ANGSD output (average pair-wise $F_{ST}$ between populations).

To compare the impact of coverage between the two approaches, we used a subset of 43 individuals that underwent an additional round of sequencing and applied the analyses on this dataset before (mean coverage of 2.47x [1.55 – 4.48]; hereafter referred to as “Low coverage set”) and after (mean coverage of 8x [3.35 – 15.24]; hereafter referred to as “High coverage set”) re-sequencing (Supplementary Figure 17).

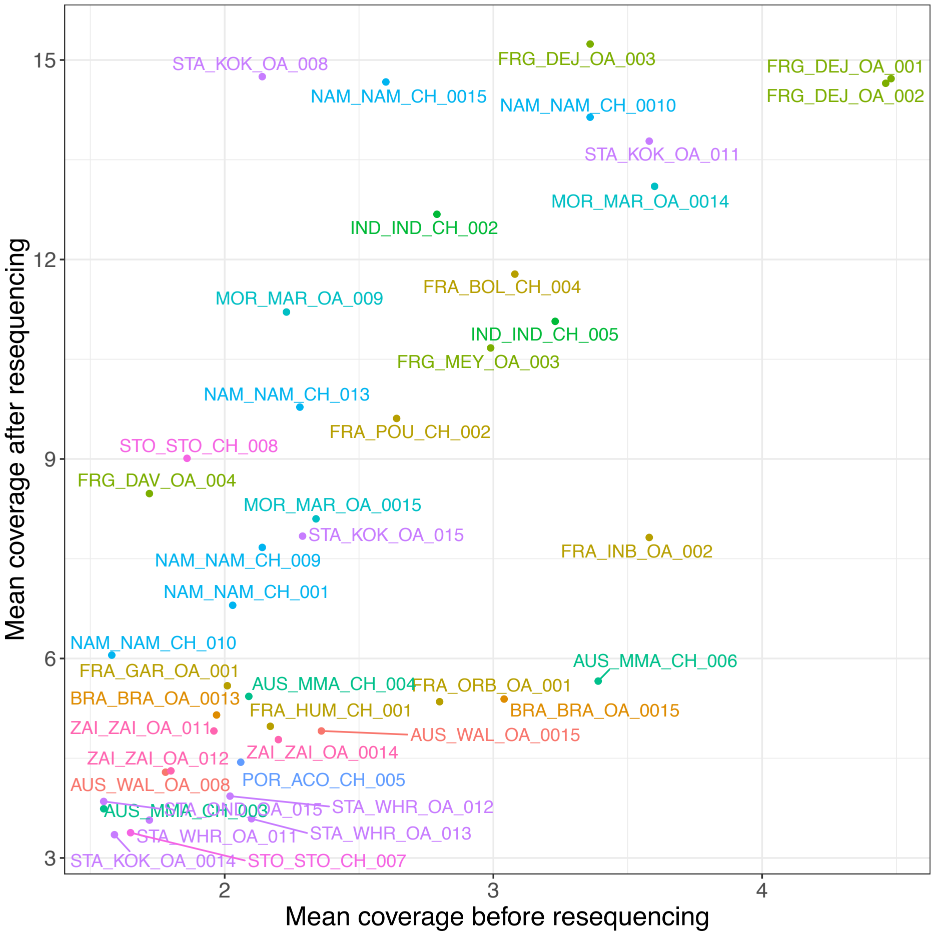


**Supplementary Figure 17. Coverage improvement for a subset of 43 individual *Haemonchus contortus* males**

Following this approach, we compared nucleotide diversity estimates and Tajima's *D* estimates in three populations exhibiting highest number of individuals with 5x mean coverage and more, i.e. Namibia (NAM), France (FRA.1) and Guadeloupe (FRG). Results (Supplementary Figure 18) demonstrated that nucleotide diversity estimates (shown for Chromosome I) were significantly biased downward in the low coverage set (-0.135%, $P < 10^{-4}$), while Tajima's *D* statistic was higher in this set (+0.028, $P < 10^{-4}$). However, distribution pattern along the considered chromosome was not altered.

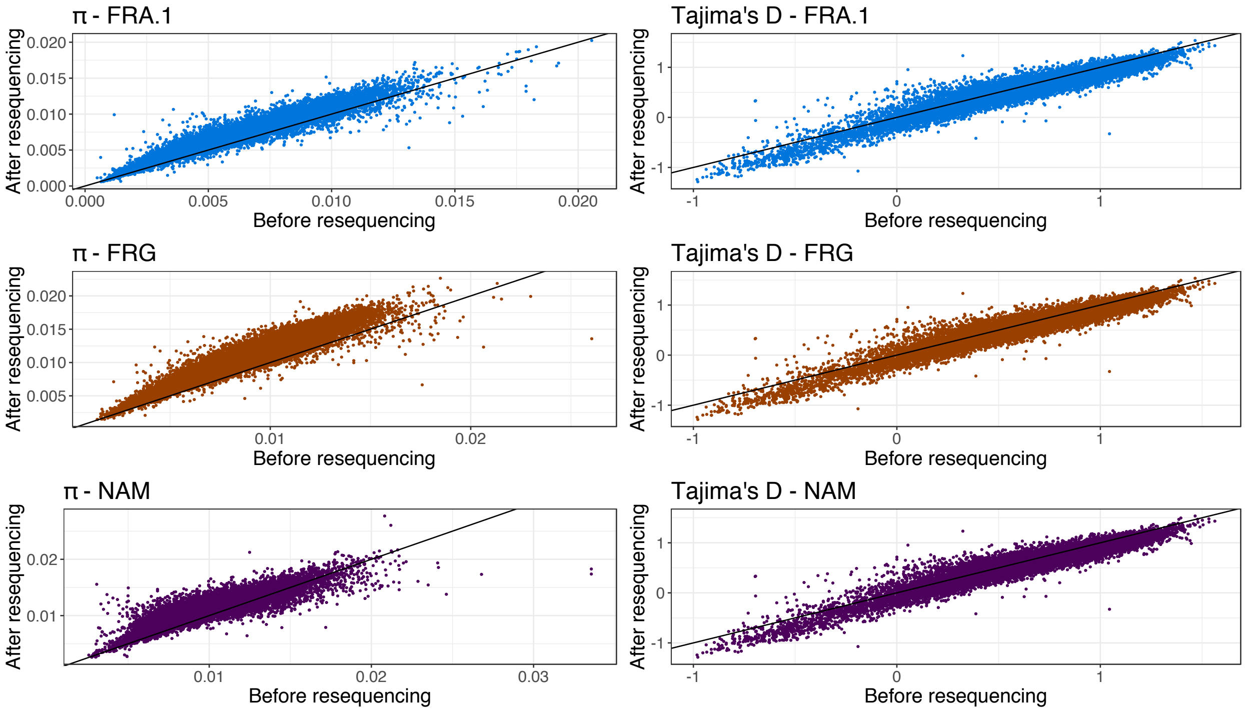


**Supplementary Figure 18. The relationship between sample coverage and population diversity estimates**

Pairwise $F_{ST}$ estimates from ANGSD were also correlated with population cross-coverage (Pearson's $r_{(136)} = 0.54$, $P<10^{-4}$), suggestive of a bias. This bias could be overcome when considering VQSR SNP calls and using the maximal value binned by 10% minor allele frequency (supplementary figure 19). In this case, correlation was not significant (Pearson's $r_{(136)} = -0.05$, $P = 0.55$).

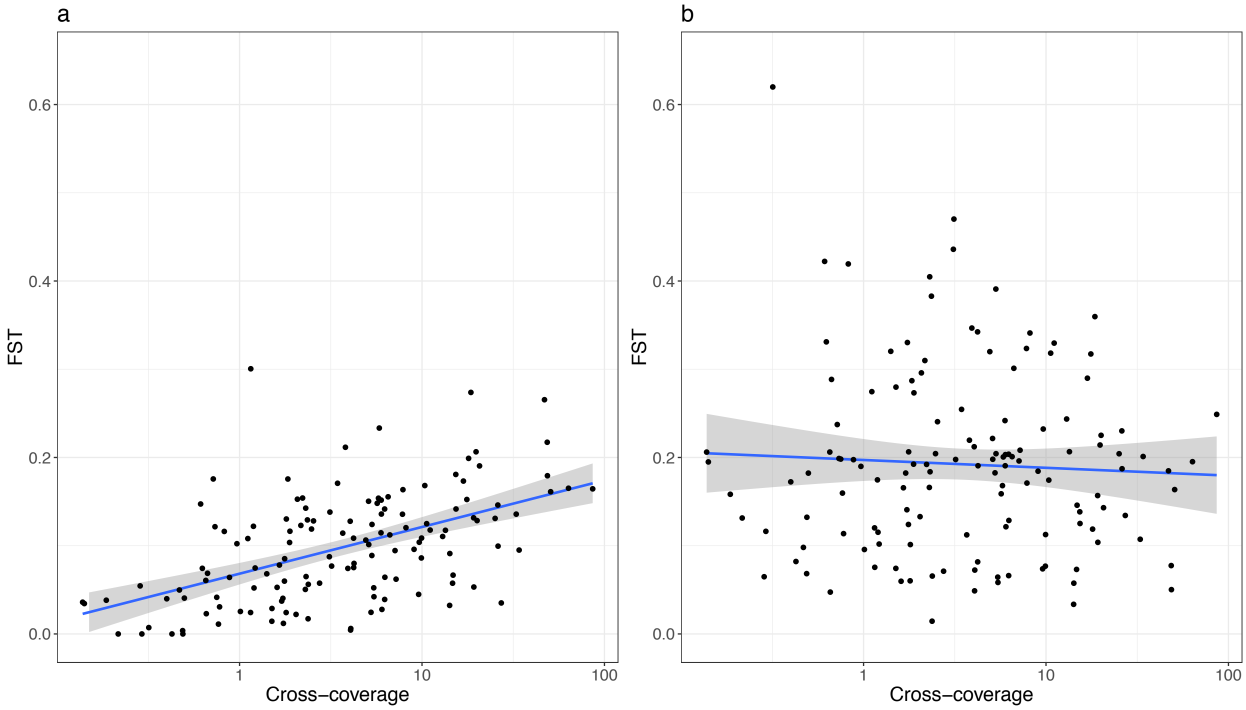


**Supplementary Figure 19. The relationship between pairwise $F_{ST}$ estimates and coverage according to the considered framework**

$F_{ST}$ values, estimated under the probabilistic framework of ANGSD (a) or using the maximal value of $F_{ST}$ binned by minor allele frequency (b) have been plotted as a function of population cross-coverage.

Pairwise divergence between individuals was also affected by coverage, as Hamming distance estimates based on VQSR SNP calls increased as cross-coverage increased (Pearson’ $r_{(49729)} = 0.31$, $P<10^{-4}$; supplementary Figure 20a). To account for this, the sole individuals exhibiting a mean depth of coverage higher than 2.5x were considered resulting in bias removal (Pearson’ $r_{(8281)} = 0.02$, $P = 0.09$; supplementary Figure 20b).

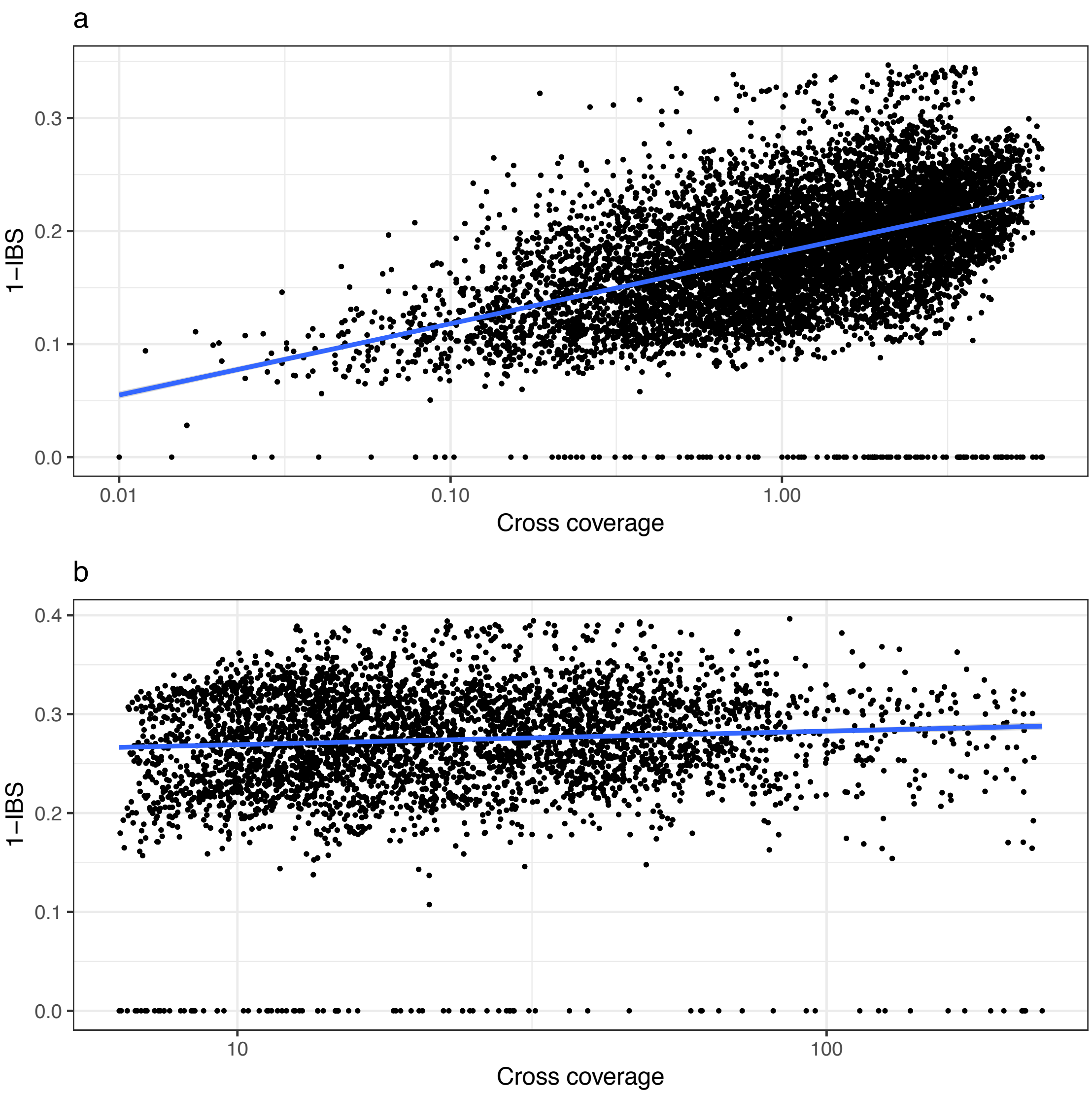


**Supplementary Figure 20. The relationship between Hamming’s distance and coverage**

Both frameworks also yielded similar population clustering. ANGSD provided more distinct clusters, especially for populations with the least coverage like Benin and the 4th French population (FRA.4). The only discrepancy was the clustering of Indonesian population among Guadeloupian samples when considering VQSR SNP calls (supplementary figure 21). The low proportion of variance explained by the first two components with ANGSD (figure 1b) remains unexplained. The first two components of a PCA on VQSR SNP explained 7.96% and 5.86% respectively.

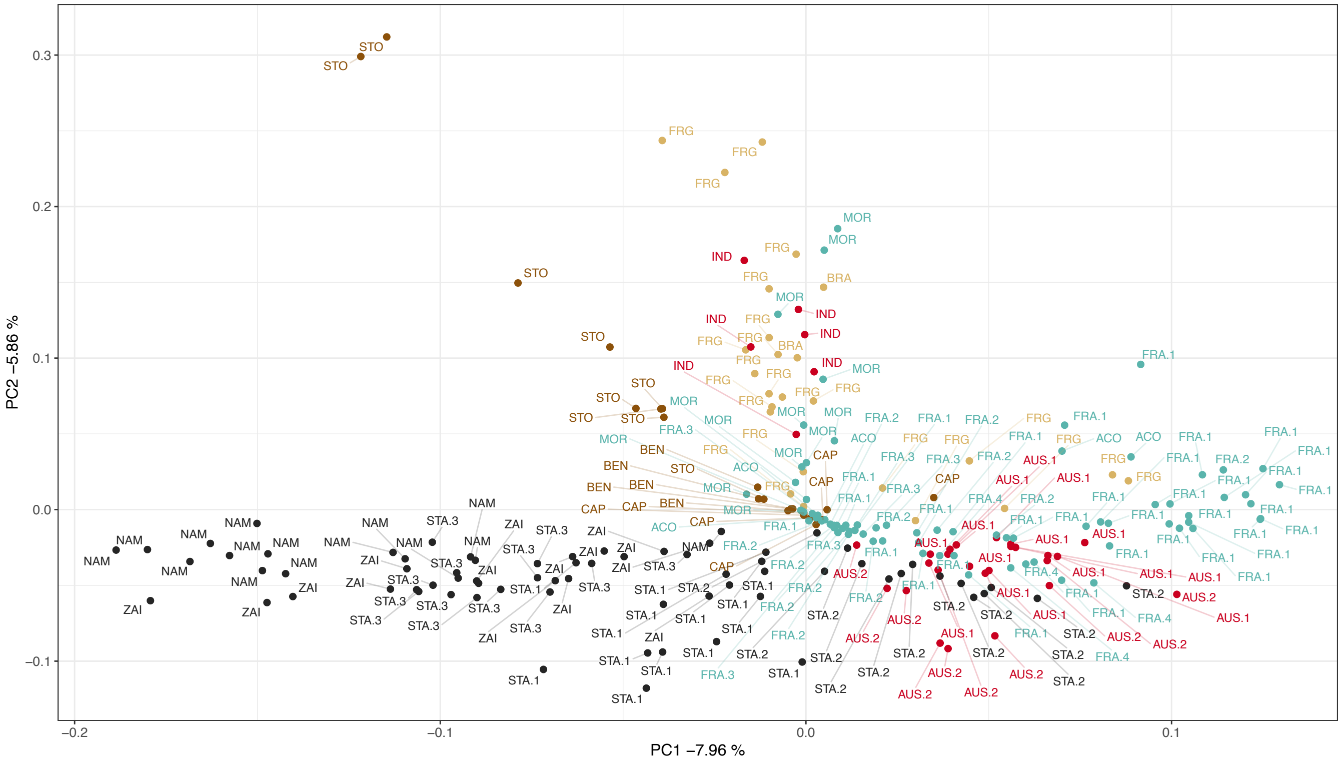


**Supplementary Figure 21. Principal component analysis based on nuclear VQSR SNP calls across 223 individuals**

Admixture analysis was robust to coverage (supplementary figure 22) as demonstrated by an analysis on the 43 samples that underwent re-sequencing. Population ancestry pattern inferred from re-sequenced samples (mean coverage of 8x [3.35 – 15.24]; right column) remained similar to that observed before re-sequencing (mean coverage of 2.47x [1.55 – 4.48]; left column).

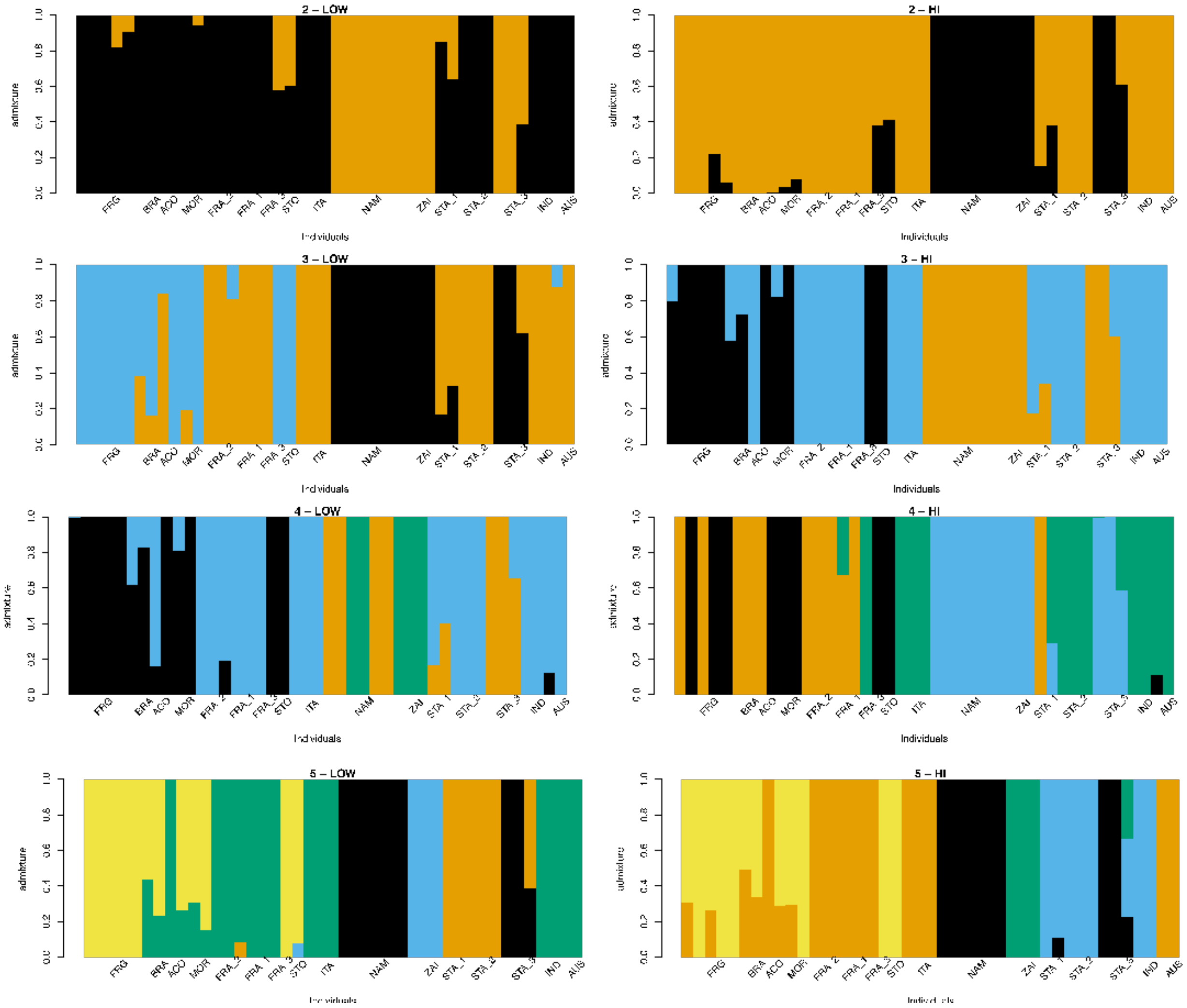


**Supplementary Figure 22. Admixture analysis run for chromosome II on the same set of 43 individuals before (left) or after (right) resequencing for K ranging from 2 to 5**

**Beagle imputation accuracy**

The topology weighting analysis implemented with TWISST and the cross-coalescent time estimation both relied on Beagle[11] imputed genotypes. To evaluate how imputation accuracy would perform on low coverage samples, Beagle was run for chromosome I on the subset of 43 re-sequenced samples, using the genotypes called on low coverage BAM files (before re-sequencing). Discordance at the individual and site levels between imputed genotypes and genotype called using data after resequencing were subsequently computed with vcftools[12] v0.1.15 using the --diff-site-discordance and --diff-indv-discordance options. At the individual level, the mean discordance was 7.2% [3.4% – 18.97%]. Discordance level was consistent throughout the chromosome (supplementary figure 23a) and two third of the sites displayed less than 10% (supplementary figure 23b) discordance between imputed genotypes and genotype called from high coverage BAM files. Discordance displayed was slightly increased for samples with higher coverage before resequencing (Supplementary Figure 23c).

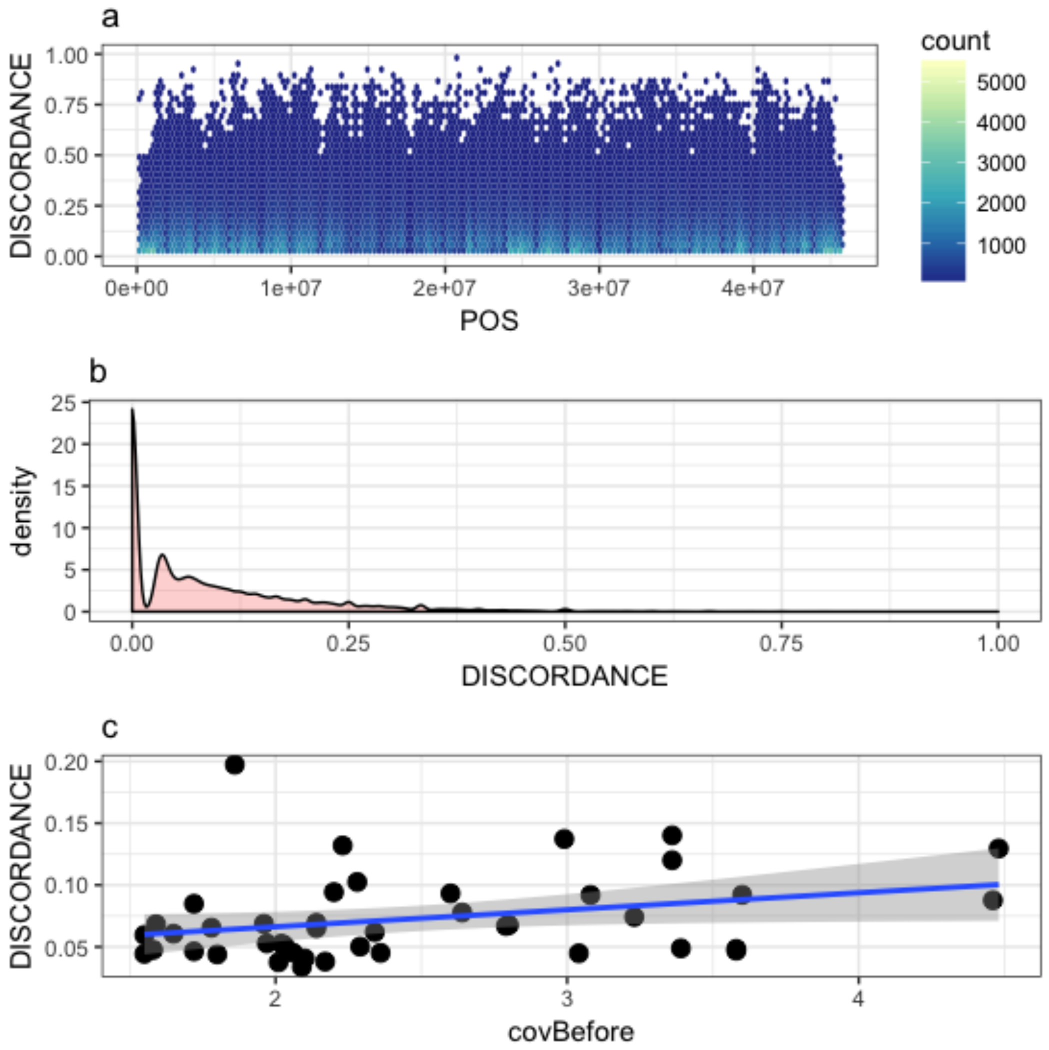


**Supplementary Figure 23. Genotype discordance after Beagle imputation**

**(a)** the density distribution of discordance (ranging between 0 and 100%) for SNP sites (binned into hexagons) along chromosome I between imputed genotypes and called genotype of the same sample with high coverage.
**(b)** the density distribution of discordance values at the SNP site level, highlighting that a majority of sites show less than 10% discordance between imputed and called genotypes with high coverage data.
**(c)** the relationship between individual coverage and the average discordance at the SNP level for chromosome I.
